# Designing Single Cell RNA-Sequencing Experiments for Learning Latent Representations

**DOI:** 10.1101/2022.07.08.499284

**Authors:** Martin Treppner, Stefan Haug, Anna Köttgen, Harald Binder

## Abstract

To investigate the complexity arising from single-cell RNA-sequencing (scRNA-seq) data, researchers increasingly resort to deep generative models, specifically variational autoencoders (VAEs), which are trained by variational inference techniques. Similar to other dimension reduction approaches, this allows encoding the inherent biological signals of gene expression data, such as pathways or gene programs, into lower-dimensional latent representations. However, the number of cells necessary to adequately uncover such latent representations is often unknown. Therefore, we propose a single-cell variational inference approach for designing experiments (scVIDE) to determine statistical power for detecting cell group structure in a lower-dimensional representation. The approach is based on a test statistic that quantifies the contribution of every single cell to the latent representation. Using a smaller scRNA-seq data set as a starting point, we generate synthetic data sets of various sizes from a fitted VAE. Employing a permutation technique for obtaining a null distribution of the test statistic, we subsequently determine the statistical power for various numbers of cells, thus guiding experimental design. We illustrate with several data sets from various sequencing protocols how researchers can use scVIDE to determine the statistical power for cell group detection within their own scRNA-seq studies. We also consider the setting of transcriptomics studies with large numbers of cells, where scVIDE can be used to determine the statistical power for sub-clustering. For this purpose, we use data from the human KPMP Kidney Cell Atlas and evaluate the power for sub-clustering of the epithelial cells contained therein. To make our approach readily accessible, we provide a comprehensive Jupyter notebook at https://github.com/MTreppner/scVIDE.jl that researchers can use to design their own experiments based on scVIDE.

## Introduction

Scientists are increasingly using experimental methods to study the transcriptome in cell populations at the single-cell level aiming to detect (sub-)types of cells, and cell type specific gene expression patterns [1]. The high complexity of the resulting data often prompts researchers to resort to unsupervised pattern discovery approaches. A class of techniques that has proven particularly useful in many such scenarios is deep learning, in which neural network approaches have been specifically adapted to the particularities of single-cell RNA-sequencing (scRNA-seq) data [2–4]. More specifically, deep generative models, such as variational autoencoders (VAEs), which learn a lower-dimensional latent representation of the cells’ transcripts have been used for visualization [5], imputation [2, 3, 6], to generate cell perturbations in-silico [7, 8], or to augment scRNA-seq data with synthetically generated data to improve robustness of downstream analyses [9].

However, there is a lack of statistical methods that could help scientists design their experiments, particularly regarding the number of cells to be sequenced, such that deep learning techniques can reliably detect potential patterns. Traditional approaches for statistical power analysis focus on somewhat less complex analysis tasks, such as differential expression analysis [10–12]. To guide the design of experiments with complex pattern discovery tasks, we propose a deep learning approach for generating synthetic cells from existing scRNA-seq data, e.g. pilot data, and subsequently perform power analyses for complex pattern discovery tasks with varying sizes of synthetic datasets.

Proper design of single-cell sequencing experiments is crucial for cell type detection, as is indicated in several review articles [13–17]. The Sajita Lab - “How many cells?” tool [18] provides a theoretical lower bound for the number of cells required to find at least *n* cells of each type. For this, the number of cell types has to be prespecified. Accordingly, the tool requires knowledge about the expected number of cell types, which is often incompatible with an exploratory clustering approach.

As noted by [19], there are already a number of methods guiding the experimental design of scRNA-seq studies when it comes to univariate endpoints like differential expression testing [10–12]. However, scientists increasingly want to answer multivariate questions, e.g., dimensionality reduction and clustering, and the design of robust studies is hindered by the lack of available methods.

Generally, many approaches for designing scRNA-seq experiments rely on simulated data, where researchers either assume linear relationships between genes [20], use kinetic models [21], simple generative models [22], or unsupervised techniques [23], which require large reference data sets. However, sample size estimation tools for learning low-dimensional representations, especially for capturing non-linear relationships, stay mostly unaddressed.

We present a method to determine the statistical power for deep learning based cell type detection with varying sample sizes. Our method is based on a deep generative modeling approach, single-cell variational inference (scVI) [2], developed to analyze scRNA-seq data. In detail, we obtain lower bounds of the log-likelihood—the probability of observing the data given the model— of scVI before and after training. Next, we compute the differences in the log-likelihood lower bounds for each cell to quantify the contribution each cell has made to the network’s training. We do the same on a data set modified by a permutation approach to obtain a null distribution for reference. More precisely, we use jackstraw sampling [24, 25], where only the values from a small subset of cells are permuted. By comparing the resulting contributions between the original and the jackstraw data, we can estimate statistical power and hence guide experimental design.

## Methods

Typically, the goal of a deep generative model (DGM), such as VAEs trained by variational inference, is to learn a joint distribution over all input variables and thereby infer an approximation to the data generating process. To model the data-generating process of the gene expression count data, we denote *x* as a vector of observed counts. Specifically, *x*^(*i*)^represents a single cell, where its components correspond to the gene expression counts of all measured genes. The task of our model is to learn the complex relationships between the genes. For example, groups of genes that exhibit routinely coordinated expression patterns could indicate underlying gene programs. To do this, DGMs use latent variables *z*, which have far fewer dimensions, e.g., only ten dimensions, than our observed gene expression space *x*. By conditioning the data-generating process on these latent variables, we can represent the joint probability distribution of *x* and *z* as:

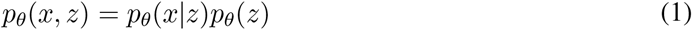

Here, *p_θ_*(*x|z*) represents the likelihood function and *p_θ_*(*z*) gives the prior probability distribution of *z* [4].

### Variational autoencoders

Variational autoencoders provide a framework that consists of two parametric models that work together to produce a lower-dimensional latent representation of the input data and, at the same time, learn how to reconstruct the input to potentially produce realistically looking synthetic data. The model that produces the latent representation is commonly referred to as the encoder, and the generative model is called a decoder. The two sets of corresponding parameters are *ϕ* and *θ*, respectively. Using stochastic gradient descent, the optimization objective can be jointly optimized for all parameters. First, we aim to approximate the posterior distribution of the latent variables *z* given the network’s input *x*, which is denoted as *q_ϕ_*(*z|x*) (and is an approximation to the true posterior *p_θ_*(*z|x*)). Next, we reconstruct the input given the latent variables using the generative model *p_θ_*(*x|z*). The marginal log-likelihood is made up of the marginal log-likelihoods for the individual data points; here cells:

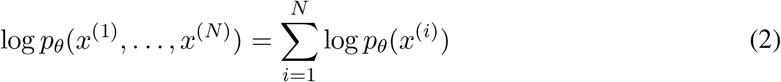

The marginal log-likelihood can then be rewritten into two interpretable terms:

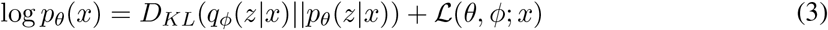

Here, the left term, *D_KL_*, represents the Kullback-Leibler (KL) divergence between the approximate posterior *q_ϕ_*(*z|x*) and the true posterior *p_θ_*(*z|x*). The second part of Equation 3 represents the evidence lower bound (ELBO), which provides a lower bound on the log-likelihood of the data:

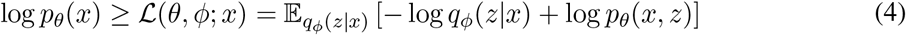

The ELBO can then, in turn, be represented as follows:

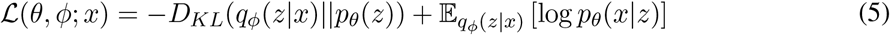

[26] [27].

For an i.i.d. data set, the ELBO objective is defined as the sum of each individual data point’s ELBO. This provides us with a lower bound about how well the model captures each specific data point or cell’s contribution [26] [27].

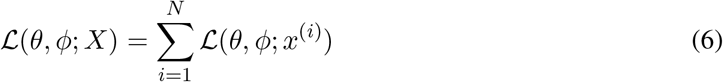

Finally, this lower bound can be optimized for both the parameters of the encoder *ϕ* and the decoder *θ*.

### Single-cell variational inference

One of the first and most common deep learning methods to learn the latent structure of scRNA-seq data is scVI, a framework based on the architecture of VAEs [2]. It encodes the scRNA-seq data into a low-dimensional latent representation in which the underlying biological signals are potentially extracted. Based on this latent representation, the network then tries to reconstruct the input data, learning to generate realistic data simultaneously.

Specifically, the variational distribution *q*(*z,l|x,s*) tries to approximate the posterior distribution of the latent variables given the data *p*(*z, l|x, s*), where *z* is a lower-dimensional vector of Gaussians, *l* represents a library-size scaling factor, *x* is a vector of observed gene expressions, and *s* encodes the batch annotation for each cell. Hence, the variational distribution is mean-field and can be written as [2]:

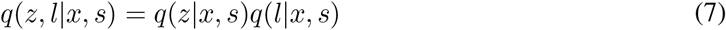

The variational lower bound can accordingly be written as:

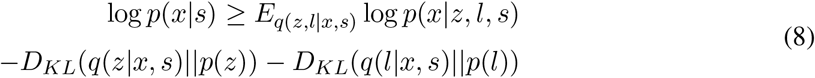

For more details on scVI, see [2].

### Cell contribution to latent representation

To investigate the contribution of each cell to the representation in the VAEs latent space, we employ a permutation method named jackstraw [24, 25]. Here, a small number *s*, say 10%, of cells are randomly selected from the cell by gene matrix, after which the expression counts for each cell are permuted between all genes (**Figure 1 A**). The permutation removes the correlation structure between genes, resulting in a null distribution for the selected cells. However, we ensure that the data set’s overall structure is preserved by limiting ourselves to a small number of observations. Hence, we can still obtain a plausible representation of the data in the latent space despite the permuted cells.

**Figure 1:**
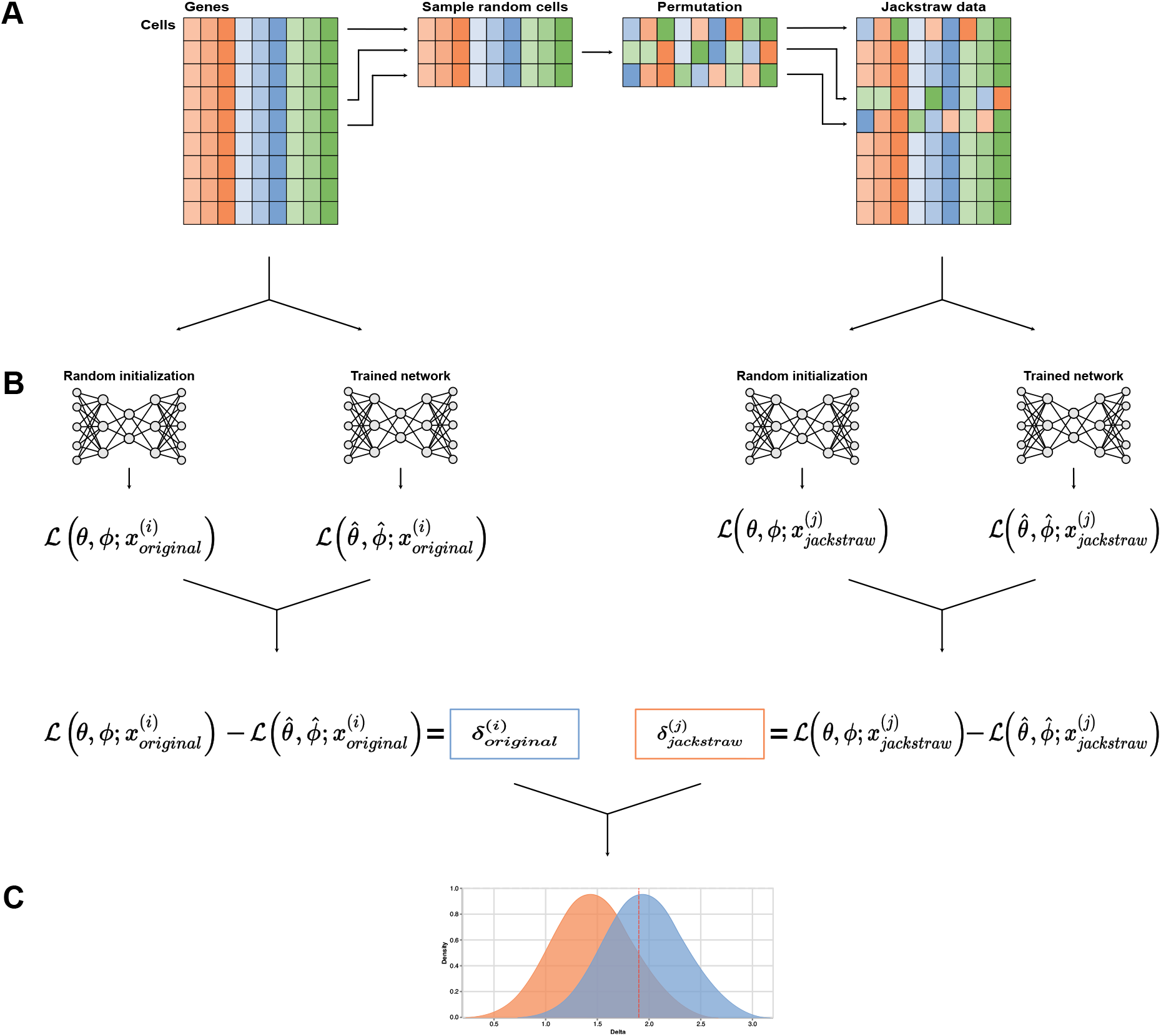
Schematic of the general steps of the scVIDE algorithm. scVIDE starts with a cell by gene count matrix from which a small number of cells (e.g., 10%) are randomly selected and counts are randomly permuted across genes. The permuted and hence uninformative cells are subsequently pasted back into the cell by gene count matrix, which we refer to as jackstraw data based on the publication by [24] (A). Next, we randomly initialize scVI for the original count matrix and the jackstraw data and compute the ELBO as a reference value for the training progress. We subsequently train scVI on the original and jackstraw matrices and compute the difference in the ELBO (B). The permutation and training on the jackstraw data are repeated *B* times to attain a sufficiently large number of cells for the null distribution. The resulting difference can subsequently be used to assess statistical power (C).

To quantify the impact of each cell on the latent representation, we consider the differences in logarithmized ELBO (see **Equation 5**) from a randomly initialized network, (θ, ϕ), with those from a trained network, 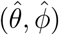. In doing so, we distinguish between a network whose cells were not permuted and several networks in which a small number of cells were permuted.

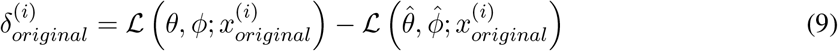

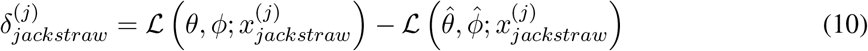

Here, 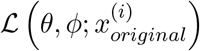 and 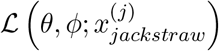 act as a reference to evaluate each cells contribution to reducing the ELBO. Additionally, 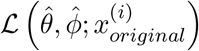 and 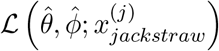 represent the estimated ELBOs after model training. Hence, we assume that permuted cells that do not introduce any additional information to the network also result in a lower ELBO reduction. Similar to the bootstrap, we repeat the jackstraw permutation *B* (*b* = 1,…, *B*) times, which means that we also have to train *B* jackstraw networks. However, since our focus is on small data regimens and training the jackstraw networks can be parallelized, the computational burden is still feasible, which we demonstrate in an exemplary runtime analysis of scVIDE using the *Wu data set* (see **Supplementary Table 3**).

### Determining statistical power

By repeated jackstraw sampling, we obtain a null distribution which we use in the next step to estimate the statistical power for learning a biologically meaningful latent representation in scVI. We use this null distribution of ELBOs across all cells to generate an empirical cumulative distribution function (eCDF). Put simply, we use the eCDF of the jackstraw ELBOs as an empirical measure and evaluate the probabilities of the ELBOs from the original data under the eCDF; i.e., assess how likely they are under the jackstraw eCDF.

The eCDF of jackstraw ELBOs 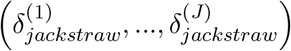 is given by:

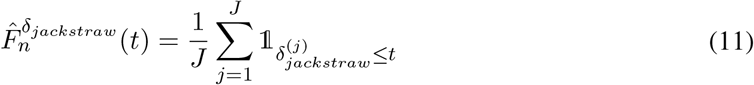

By evaluating the original data ELBOs on the complementary CDF of **Equation 11** we obtain one-sided p-values while treating observed values of *δ_original_* as test statistics:

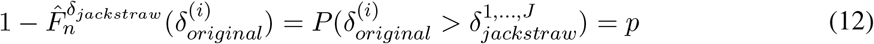

Doing this, we infer how often ELBO differences derived from informative cells exceed the ones derived from uninformative jackstraw cells, which can be formulated in the following hypotheses:

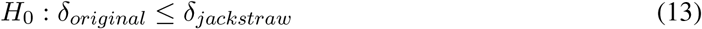

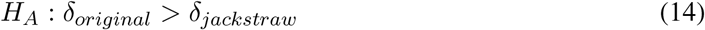

Thus, we test whether training on the original data represented as 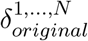 significantly decreases the ELBO compared to training on uninformative data, which we represent as 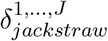.

With the above approach, we can determine the statistical power for a given number of cells. To assess the statistical power for experiments whose number of cells exceeds that of the pilot dataset, we can use the properties of VAEs and generate synthetic data. Using the generated data that exceeds the size of the original dataset, we can then re-apply the algorithm to obtain the statistical power for the enlarged dataset. Finally, we can continue this procedure until achieving adequate statistical power.

### Experimental setup

To test scVIDE, we used the following setup. First, we trained the algorithm on varying sizes of the original small pilot data sets n = (64, 128, 192, 256, 320, 384, 768, 1152, 1536). Here, we arrived at the small pilot data sets by randomly subsampling cells from the original larger data. Note that when we refer to pilot data sets in the following, we mean smaller subsamples of the original data that are assumed to represent pilot data (**Figure 2 A**). Based on the trained models, we then generated synthetic data, first in the size of the input pilot data set and then by synthetically increasing the sample size 2-fold, 3-fold, 4-fold, and 5-fold. For example, after training the model on 384 cells, we would generate 384, 768, 1152, 1536, and 1920 synthetic cells (**Figure 2 B**). Additionally, for each pilot data set, we sample 64 cells and generate the same amount of synthetic cells using scVI. Next, we train scVI on the 64 cells to estimate each cell’s ELBO based on the correlation structure provided by the synthetic data. To create the null distribution of ELBO estimates, we would use jackstraw sampling on a small fraction, say ten cells, of the 64 cells and train scVI on the permuted data set. Repeating this procedure 100 times, we could therefore estimate the ELBO of 1000 uninformative cells. We can then use the resulting distribution of ELBO estimates—stemming from uninformative cells— as a baseline to quantify the contribution of a potentially informative cell to the ELBO. Hence, we always compare ELBOs from the larger data sets to that of the small 64-cell pilot data, limiting computational complexity. We only apply scVIDE to synthetic data because, in reality, we would have only a small pilot data set at hand. Synthetic data can be seen as a noisier version of the original data. Hence, we expect statistical power derived from synthetic data to give a lower bound for the original data. Next, we applied scVIDE to each synthetic data set and estimated the statistical power for each sample size. For the jackstraw sampling, we have set the proportion of permuted cells to s = 10% and trained B = 100 networks based on the permuted data. In a sensitivity analysis, we show that our model is robust to the choice of the hyperparameter (see **Supplementary Figure 2**).

**Figure 2:**
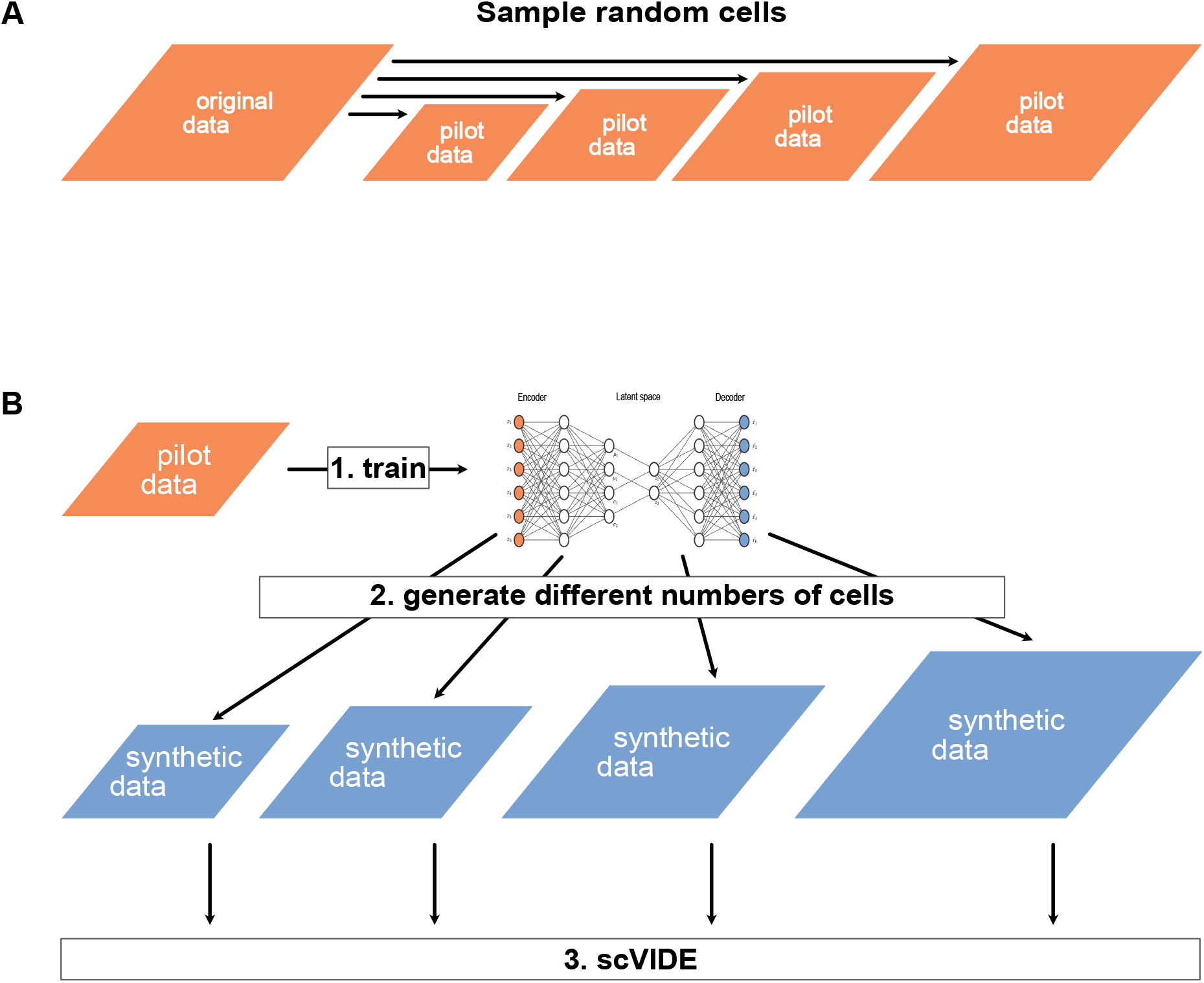
Experimental set up for testing the performance of scVIDE. We generate pilot data sets by randomly sampling from a large data set (A). Next, scVI can be trained on the small pilot data sets, after which we can use the resulting model to generate synthetic data in varying sizes. We apply scVIDE to synthetic data of size n = (64, 128, 192, 256, 320, 384, 768, 1152, 1536) (B). The resulting statistical power for different sample sizes can then be used to plan future experiments.

## Results

We applied scVIDE to five data sets representing different experimental protocols and tissues. Specifically, we use a data set from the cortex of mouse brains [28], which we refer to as *Tasic data* throughout this work. In addition, we use a data set of peripheral blood mononuclear cells (PBMCs) from a healthy donor [29], where we refer to this data set as *PBMC3k*. We additionally examine a Smart-seq2 data set (*Nestorowa data*). This data set consists of mouse hematopoietic stem cells (HSC) [30].

For these three data sets, we have computed a theoretical bound based on the cumulative distribution function (CDF) of the negative binomial distribution as proposed by [18]. This bound estimates the number of cells that need to be sampled to detect a minimum amount of cells from each cell type. Since the approach assumes that the population of cells is dominated by one cell type, detection of less abundant cell types will be more challenging, which leads to higher estimates for the necessary number of cells. Also, this theoretical approach does not take technical or biological factors into account [21]. Therefore, we expect scVIDE to provide higher statistical power for smaller sample sizes, and we treat the CDF-derived sample size estimates as an upper bound. We derived the input parameters for the expected number of cell types, the minimum fraction of the rarest cell type, and the minimum of the desired number of cells per type from the original publications (**Supplementary Table 2**).

Furthermore, we use two kidney data sets. First, to investigate the effects of different numbers of genes on learned latent representations, we consider a single-nucleus RNA sequencing (snRNA-seq) data set from the healthy mouse kidney, which we call *Wu data set* [31]. Also, we use snRNA-seq data from the Kidney Precision Medicine Project (KPMP) [32] to demonstrate the properties of scVIDE for power calculations in sub-clustering tasks. We refer to this data set as *KPMP data*. All data sets are publicly available.

### scVIDE provides estimates of statistical power for learning latent representations

First, we ran scVIDE on the *Tasic data*, which was generated using the SMARTer protocol. After pre-processing, the cell by gene count matrix consisted of 1525 cells and 105 genes. We randomly subsampled ten pilot data sets of each sample size (64, 128, 192, 256, 320, 384, 768, 1152, 1536 cells) and trained scVI on the resulting samples. Next, by taking advantage of the generative properties of scVI, we simulate synthetic data and increase the sample size up to fivefold. Running scVIDE on each of the ten synthetic data sets of each sample size, we observe a low statistical power. For pilot data sets of sizes 64, 192, and 384 cells, we generally stay below a statistical power of 0.5. Increasing the number of cells to approximately 1500 improves statistical power up to around 0.6. However, after further expanding the sample size, statistical power saturates. This might be related to the low number of genes in the original dataset, which only allows uncovering a limited amount of latent structure (**Figure 3 A**). The difference between null and alternative distributions of δ-values and the resulting differences in statistical power can also be illustrated using the eCDFs (**Supplementary Figure 1**).

**Figure 3:**
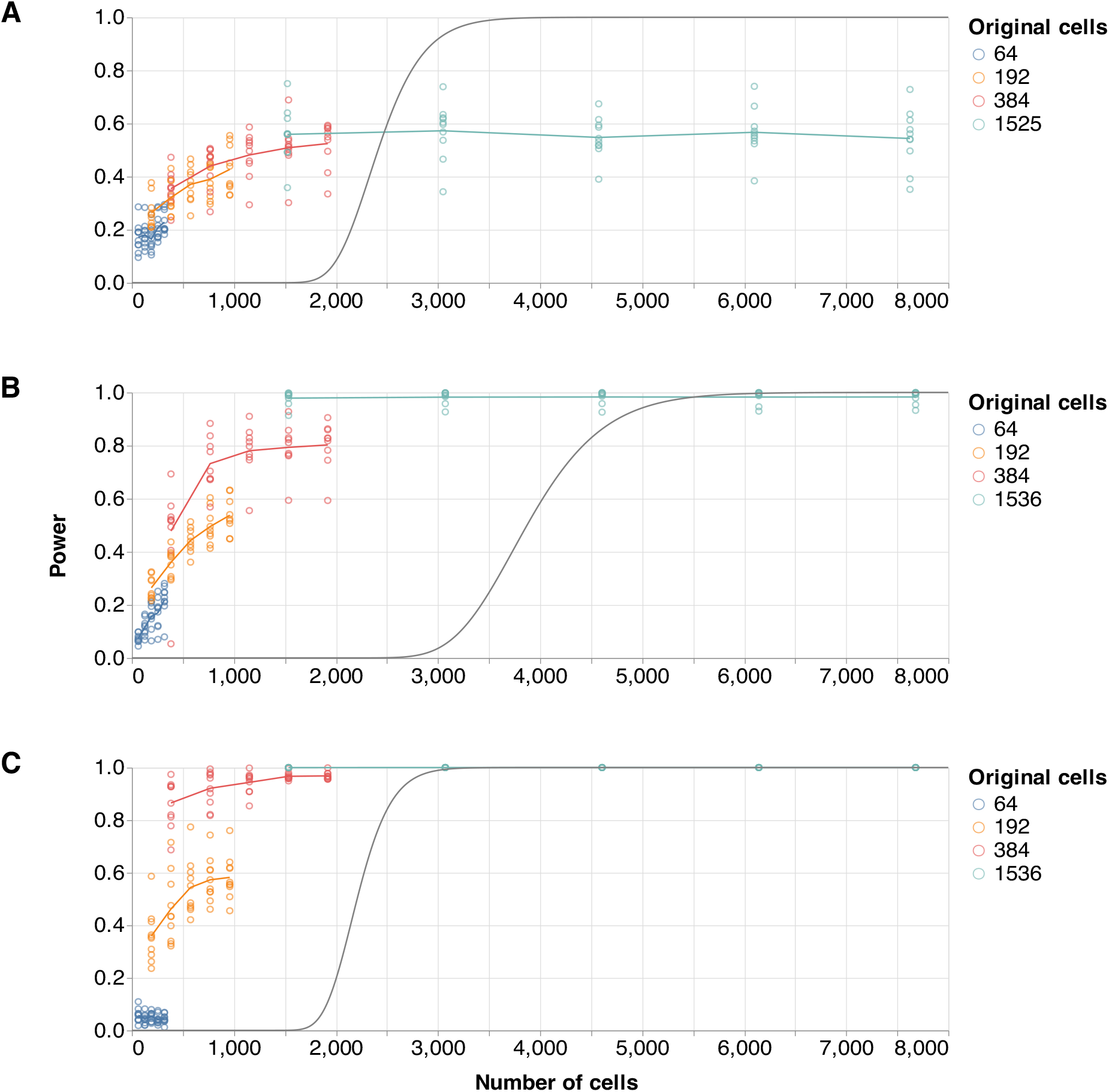
Sample size estimation using scVIDE. scVIDE can be used to estimate the sample size for scRNA-seq experiments based on smaller pilot data sets. Shown are the statistical power for ten training iterations (circles) and the respective means (lines) for varying numbers of cells. The colors indicate different sizes for pilot data sets that scVIDE was trained on, after which we additionally estimated statistical power for a 2-fold, 3-fold, 4-fold, and 5-fold increase in sample size by adding synthetic data generated from the model. We apply scVIDE to the Tasic data [28] (A), to the PBMC3k data [29] (B), and to the Nestorowa data [30] (C). The gray line indicates a theoretical lower bound based on the negative binomial CDF as detailed in [18].

Using a data set of peripheral blood mononuclear cells (PBMCs) from a healthy donor [29], we investigate statistical power for a data set with higher dimensionality. The data set consists of 2638 cells and 750 most highly variable genes after pre-processing and was generated using 10X Genomics protocols. Although statistical power is low for 64 and 192 cells, using a pilot data set of 384 cells is sufficient for planning larger experiments that reach a power of more than 80%. More specifically, statistical power reaches at least 80% at around 1500 cells and saturates close to 100% for higher cell numbers (**Figure 3 B**). This indicates that sequencing 2000 cells might have been enough to uncover latent structure in the data to a sufficient degree.

To evaluate differences in sequencing protocols, particularly between sparse and less sparse protocols, we additionally examined a Smart-seq2 data set (*Nestorowa data*). This data set consists of mouse hematopoietic stem cells (HSC) [30] with 1656 cells and 384 most highly variable genes left after pre-processing. Using the Smart-seq2 data set, we reach a statistical power of 80% at around 800 cells, which is a substantially smaller sample size than for *PBMC3k* and might be explained by differences in sparsity or differences in biological variation among cells (**Figure 3 C**).

When applying scVIDE to the three data sets, we see that the power curves are generally concave, meaning that power increases with increasing sample size and a constant number of genes, indicating that we can use synthetic data to plan larger experiments. As expected, we can more effectively plan future experiments with larger pilot data sets because the extrapolation ability depends on synthetic data quality, which in turn depends on the size of the pilot data.

Using an exemplary pilot data set, we can examine the jackstraw samples used by scVIDE and visualize them in two-dimensional space. When we visually inspect the two-dimensional UMAP representations [33] of the learned latent variables, we can confirm that the visual separability of the underlying structure in each data set which can be extracted correlates with the statistical power suggested by scVIDE. For small sample sizes, jackstraw samples seem to be difficult to distinguish from the original cells. In contrast, the UMAP representation for 1536 cells already suggests a clear separation between jackstraw and original samples. However, some of the original cells (orange) still seem to intermingle with the jackstraw samples even for the larger sample size (**Figure 4**).

**Figure 4:**
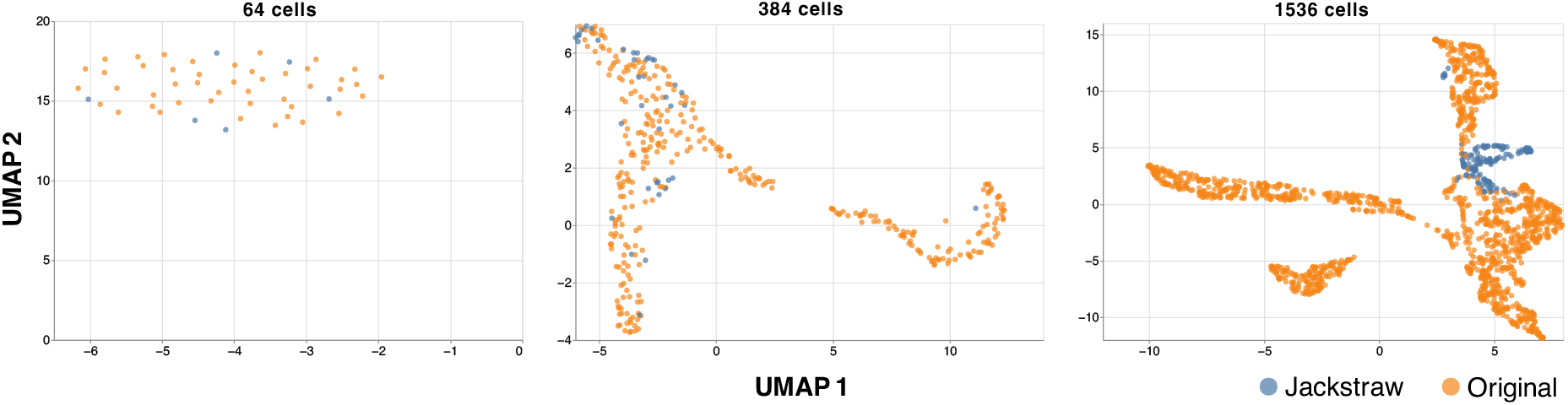
Latent representations of an exemplary data set for varying sample sizes. Using the ten-dimensional latent representation from scVI and subsequently applying UMAP for a two-dimensional representation, we can visually inspect the ability to disentangle jackstraw samples (blue) from original samples (orange). Due to the jackstraw permutation applied to the data, the UMAP representations should position jackstraw samples in a distinct location compared to non-permuted samples. The data shown comes from the *PBMC3k* data set [29].

Additionally, we find that including too many genes might impair the quality of latent representations for two reasons (**Figure 5**). First, the cell-to-gene ratio should not be too small, as this can lead to underfitting, resulting in a high bias and unstable training [2]. Underfitting can therefore be particularly problematic when generating synthetic data for larger experiments. Secondly, adding more genes from the list of the most highly variable genes simultaneously adds genes with a lower signal-to-noise ratio. This potentially leads to a noisy latent representation, which is reflected in a decrease in statistical power with an increasing number of genes. Since sample size planning is often based on small pilot data sets and the performance of scVI also decreases as the cell-to-gene ratio decreases

**Figure 5:**
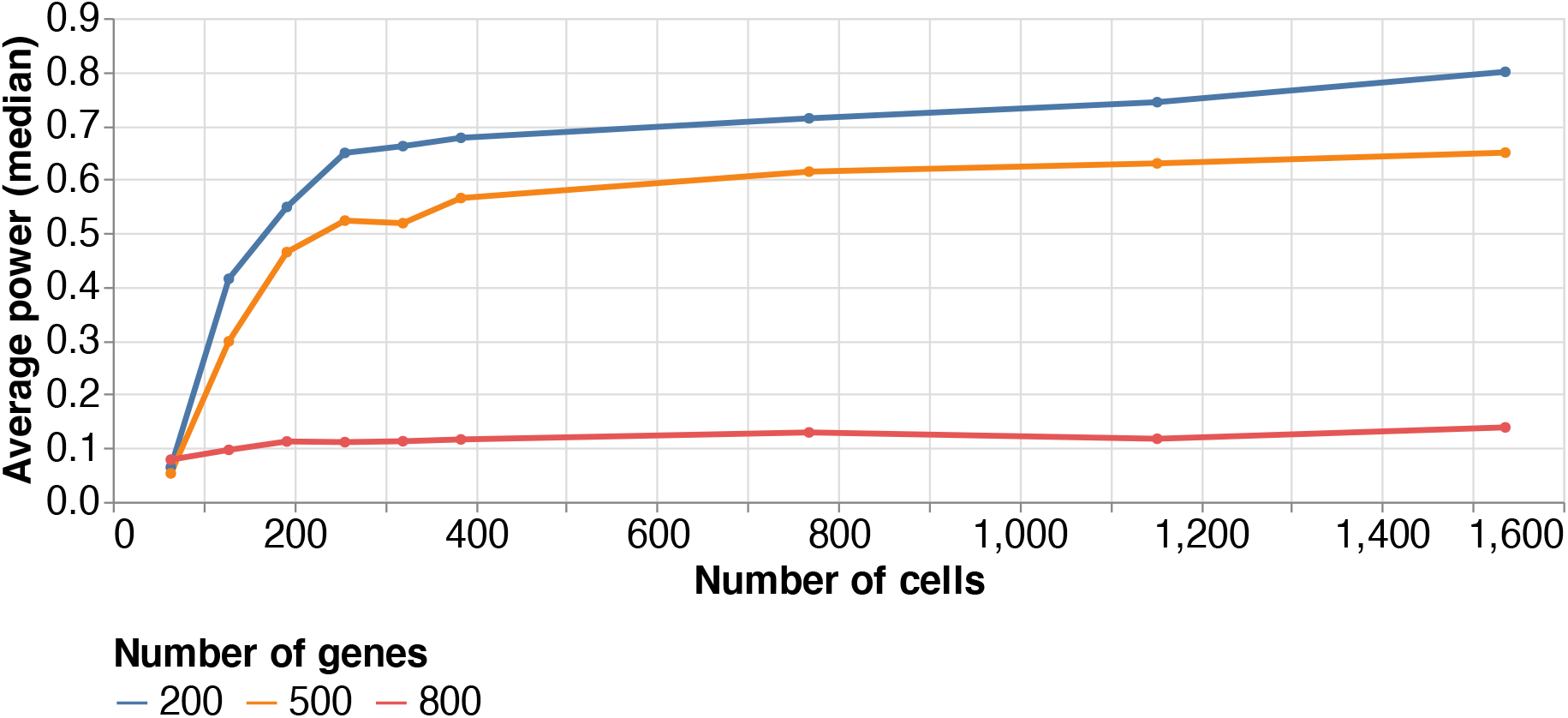
Statistical power of latent representations for varying numbers of genes. We can use scVIDE to examine statistical power for latent representations with different numbers of genes. Using the Wu *data set*, we quantify the effect of gene numbers for varying sample sizes where we investigate 200 (blue), 500 (orange), and 800 (red) most highly variable genes. The lines indicate the median statistical power over 10 replicates with different subsamples.

[2], we observe considerable losses in statistical power at 800 genes. Consequently, when planning future studies, researchers should limit themselves to genes that are relevant for their experiments to avoid unnecessarily reducing the statistical power of the latent representation. However, planning with correspondingly high sample size is inevitable if one is interested in genes with a low signal-to-noise ratio.

### scVIDE can be used to determine power for sub-cluster analyses

In addition to the typical considerations of experimental design, power calculations are also highly relevant in other scenarios. A frequently recurring issue in large cell atlas projects, such as the Human Cell Atlas [34], or the mouse brain atlas [35], is sub-clustering. In such atlas projects, cells are systematically assigned to specific cell types or cell states. The large data sets offer a high degree of resolution, i.e., larger cell clusters are split into smaller groups by sub-clustering to investigate them more closely. Using scVIDE, we can determine the possible degree of sub-clustering by estimating statistical power for models trained only on the smaller sub-clusters. By doing this, we can define limits for sub-clustering beyond which researchers lose too much statistical power to examine the individual clusters more in-depth.

To mimic a sub-clustering situation, we randomly draw N = (1000, 2500, 5000, 10000) cells from the full *KPMP data*, which included 5526 highly variable genes after pre-processing. We assume that these sampled data sets represent our entire cell population. Next, we extract only the cells that have been labeled as epithelial cells in [32] from each of the sampled data sets. The resulting epithelial cell clusters contain n = (738, 1885, 3768, 7540) cells, which we will use to estimate the statistical power for sub-clustering. In [32], the authors show that the epithelial cells can be divided into Proximal Tubule Cells (PT), Podocytes (POD) and 8 other distinct celltypes.

Applying scVIDE to the epithelial cell clusters while varying the size of the data set shows that a larger number of cells still leads to improved latent representations and thus to higher statistical power. However, the statistical power remains comparatively low even with 7540 cells. We observe that with 7540 cells, statistical power is large enough to separate POD cells but not sufficiently large to form an easily distinguishable cluster of PT cells. However, the separation of the PT cells becomes clearer as the number of cells increases (**Figure 6**). This is to be expected since cells within a cluster, in this case, the epithelial cells, are by definition more homogeneous. We could interpret the homogeneity as a small effect size similar to classical sample size calculations. Consequently, scientists generally need more cells for sub-clustering than would be the case for rather heterogeneous cell groups. We have observed a similar effect for immune cells (**Figure 3**).

**Figure 6:**
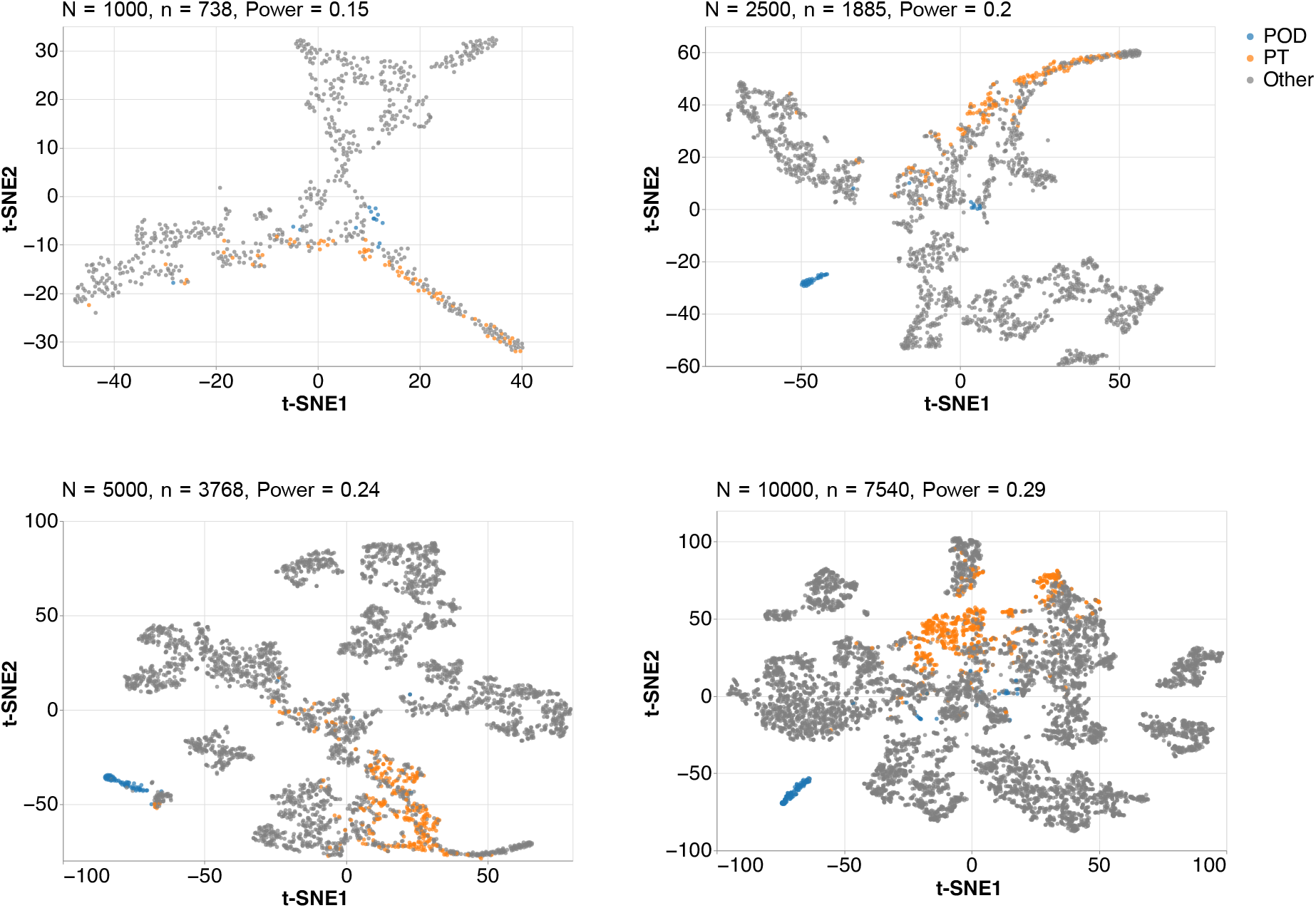
Latent representations of sub-clustering epithelial cells with varying samples sizes. Using samples of size N = (1000, 2500, 5000, 10000, 50000) from the KPMP data set [32], we extract epithelial cells and apply scVIDE to each of the resulting immune cell clusters to examine statistical power for sub-clustering. We show t-SNE representations of the lower-dimensional latent space of scVIDE and give the corresponding power estimates for each sample size n = (738, 1885, 3768, 7540, 37546) of the epithelial cells. The cell type labels are Podocytes (POD) and Proximal Tubule Cells (PT).

## Discussion

We have presented an algorithm (scVIDE) to determine the necessary number of cells for detecting potentially complex patterns in scRNA-seq datasets using deep generative models. More specifically, we adapted the existing variational autoencoder framework scVI [2], so that it allows us to estimate statistical power for pattern detection while accounting for technical factors such as batch effects. We do so by using a permutation based approach that determines each cell’s contribution to the training of scVI. Additionally, we have shown how synthetic data generated from scVI can be used to simulate and plan large-scale experiments that exceed the size of the pilot data. We also demonstrated how scVIDE can be used to determine statistical power for sub-clustering, as is often done in cell atlas projects. Our permutation framework comes with the advantage of being applicable to any model that provides an estimate of the likelihood for each cell. Hence, extending scVIDE to deep Boltzmann machines (DBMs), which have been adapted to scRNA-seq data [36], could be useful because it was previously shown that DBMs could learn from smaller data sets compared to other deep generative models [37]. Additionally, DBMs provide exact estimates of the log-likelihood, whereas VAEs only provide a lower bound, which could result in more precise statistical power estimates.

In addition to scRNA-seq experiments, scientists are now increasingly using methods that can measure multiple modalities, such as gene transcription and chromatin accessibility, in single cells. Many of the DGMs used to learn joint representations of such data are based on VAEs in which the ELBOs of the independently learned lower-dimensional latent representations of the different modalities are linked together by various mathematical operations [38]. Since this leads to a joint ELBO, as implemented in scMM [39] or Cobolt [40], scVIDE could potentially be extended to the analysis of multi-omics data sets. This is particularly interesting since joint profiling protocols currently have comparatively low throughput and, therefore, often have to cope with a smaller number of cells.

Intriguingly, we find that using a pilot data set as small as 64 cells seems sufficient for generating a reasonable null distribution that we can then use to evaluate statistical power for potentially larger data sets. An additional advantage of using real data as a starting point is that we do not assume any prior knowledge, for example, about the number of potential cell types. This allows for more generally planning experiments that aim at investigating the a priori unknown cell type composition of samples.

The overall aim of many sequencing experiments is to develop targeted treatments that are not only customized to individual patients but also to specific cell states or subsets of potentially pathological cells. These are typically detected and identified by learning lower-dimensional latent representations. We therefore expect that appropriate methods like scVIDE which can be used to design scRNA-seq studies with the necessary number of cells will be highly relevant in the future.

## Funding

M.T. was supported by the Deutsche Forschungsgemeinschaft (DFG, German Research Foundation) — 322977937/GRK2344 and DFG - Project-ID 431984000 - SFB 1453. S.H., A.K., and H.B. were supported by the Deutsche Forschungsgemeinschaft (DFG, German Research Foundation) — Project-ID 431984000 - SFB 1453.

## Availability of data and materials

The *Tasic data set* is available at https://www.ncbi.nlm.nih.gov/geo/query/acc.cgi?acc=GSE71585. The *PBMC3k data set* is available at http://support.10xgenomics.com/single-cell/datasets. The *Nestorowa data set* is available at https://www.ncbi.nlm.nih.gov/geo/query/acc.cgi?acc=GSE81682. The *Wu data set* is available at https://www.ncbi.nlm.nih.gov/geo/query/acc.cgi?acc=GSE119531, and the *KPMP data set* is available at https://qa-atlas.kpmp.org/repository/. Additionally, all analysis scripts, a comprehensive Jupyter notebook, and the scVIDE codebase are available at https://github.com/MTreppner/scVIDE.jl.

## Competing interests

The authors declare no competing interests.

## Authors’ contributions

M.T. and H.B. conceived the methods. M.T. conducted the analyses, implementations and wrote the manuscript. All authors read and approved the final manuscript.

## Supplementary Material

**Supplementary Table 1:** Hyperparameters

|  | PBMC3k | Tasic | Nestorowa | Wu | KPMP |
| --- | --- | --- | --- | --- | --- |
| Learningrate | 0.0001 | 0.00001 | 0.0001 | 0.0001 | 0.0001 |
| Hidden layers | 3 | 2 | 2 | 1 | 1 |
| Epochs | 500 | 200 | 500 | 1000 | 100 |
| Latent dimensions | 10 | 2 | 10 | 5 | 6 |

**Supplementary Table 2:** Satija lab - “How many cells?” input parameters

|  | PBMC3k | Tasic | Nestorowa |
| --- | --- | --- | --- |
| Assumed number of cell types | 15 | 10 | 9 |
| Minimum fraction (of rarest cell type) | 0.0066 | 0.0042 | 0.012 |
| Minimum desired cells per type | 10 | 11 | 20 |

**Supplementary Table 3:** Runtime analysis for an examplary run of scVIDE with sample sizes augmented by synthetically generated data using the *Wu data set* [31] with epochs = 200, genes = 500, and B = 50. Times are indicated in seconds and relate to an 2,3 GHz 8-Core Intel Core i9 with 32 GB 2667 MHz DDR4.

| scVIDE runtime analysis |  |  |
| --- | --- | --- |
| Number of cells | time | %tot |
| 64 → 1536 → 3072 → 4608 → 6144 → 7680 | 2826s | 41.9% |
| 64 → 1152 → 2304 → 3456 → 4608 → 5760 | 1061s | 15.7% |
| 64 → 768 → 1536 → 2304 → 3072 → 3840 | 780s | 11.6% |
| 64 → 384 → 768 → 1152 → 1536 → 1920 | 477s | 7.1% |
| 64 → 320 → 640 → 960 → 1280 → 1600 | 406s | 6.0% |
| 64 → 256 → 512 → 768 → 1024 → 1280 | 392s | 5.8% |
| 64 → 192 → 384 → 576 → 768 → 960 | 327s | 4.8% |
| 64 → 128 → 256 → 384 → 512 → 640 | 270s | 4.0% |
| 64 → 128 → 192 → 256 → 320 | 209s | 3.1% |

**Supplementary Figure 1:**
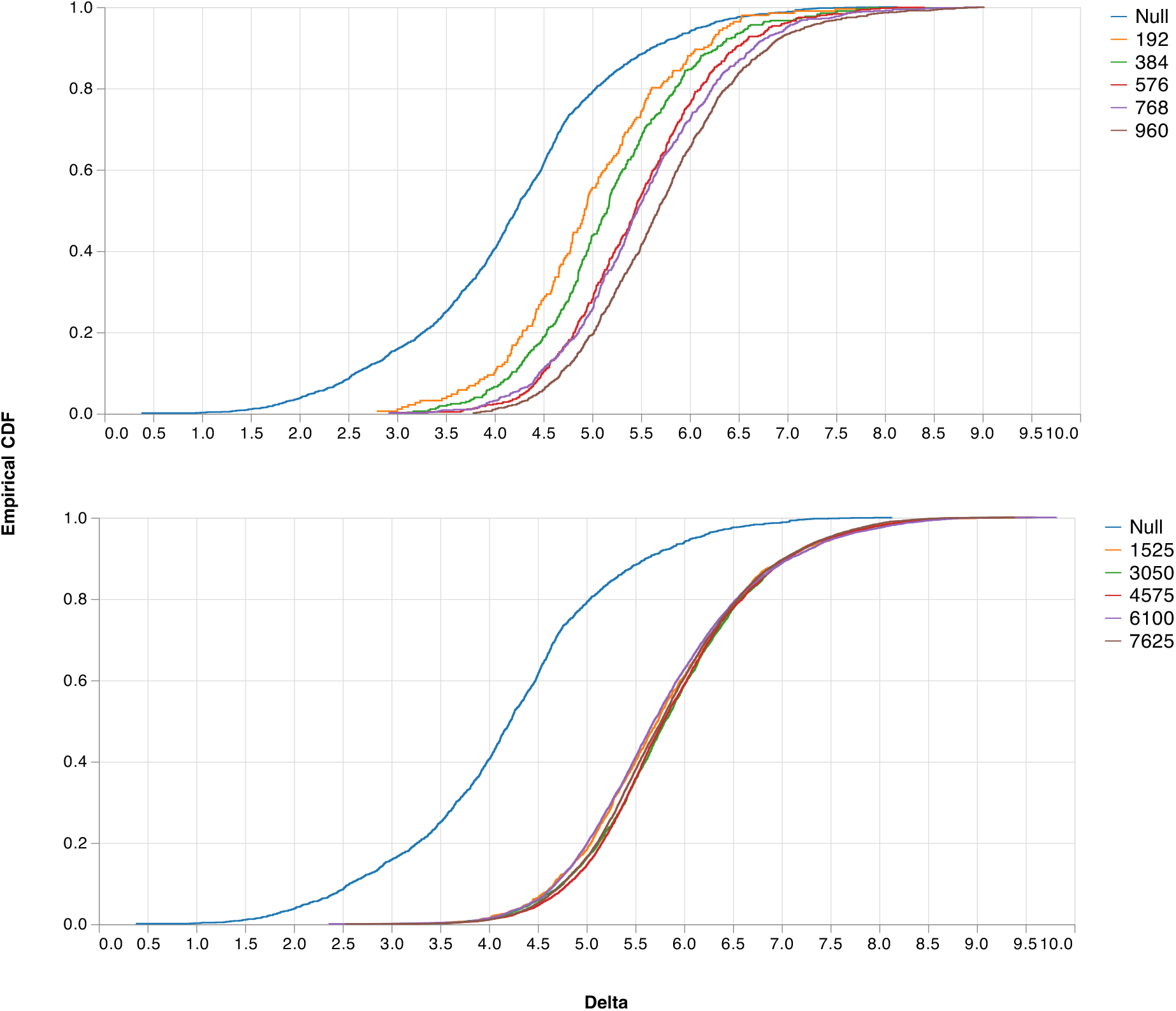
Comparing eCDFs of the null distribution with a varying number of synthetic cells. The shift in 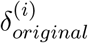 values for a small pilot data set of 192 cells compared to the null distribution of 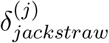 values (top). Furthermore, we show the eCDF 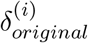 values for a pilot data set size of 1525 cells and the corresponding null distribution of 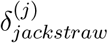 values based on 64 cells (bottom). Cell numbers larger than the pilot data indicate synthetically augmented data sets.

**Supplementary Figure 2:**
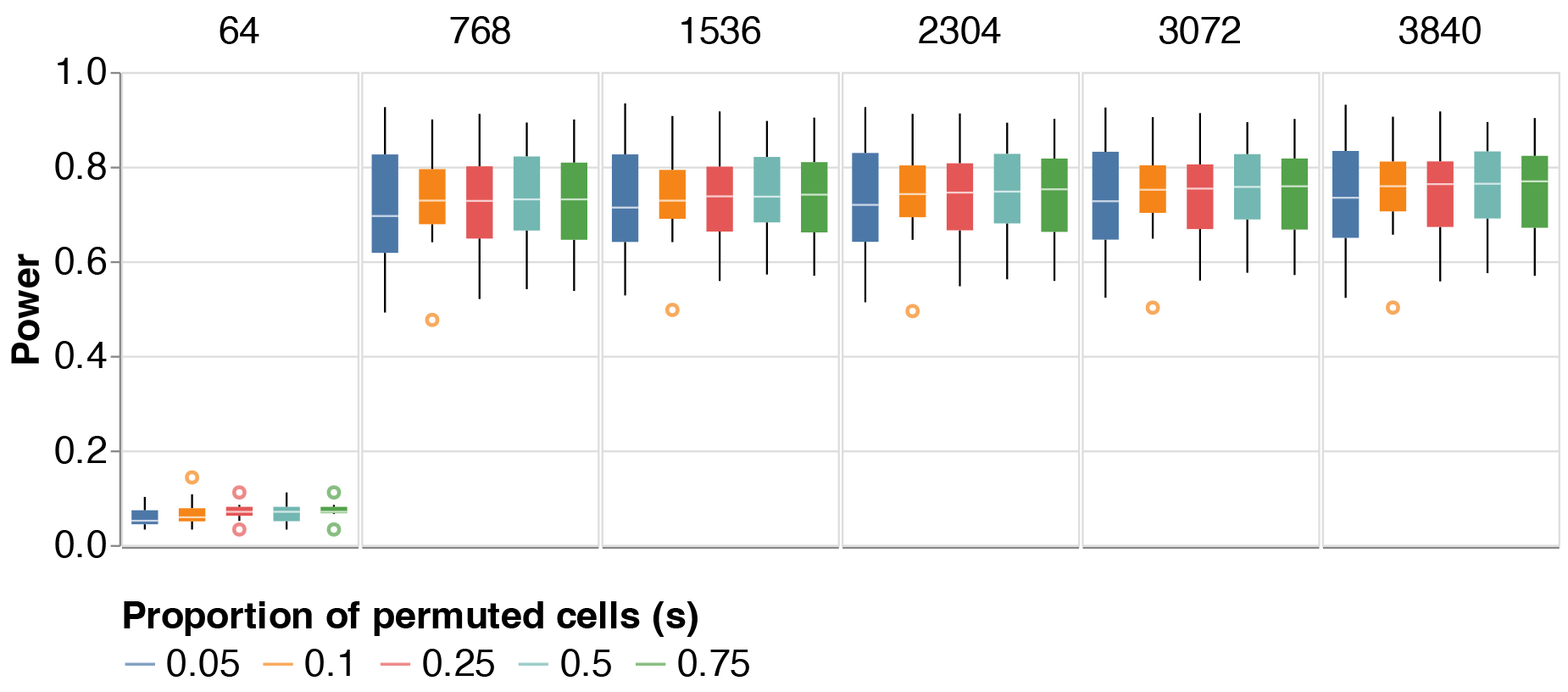
Sensitivity analysis of hyperparameter *s*. Power for varying proportions of permuted cells (s) and different numbers of cells for the *Wu data set* with 200 genes.

**Supplementary Figure 3:**
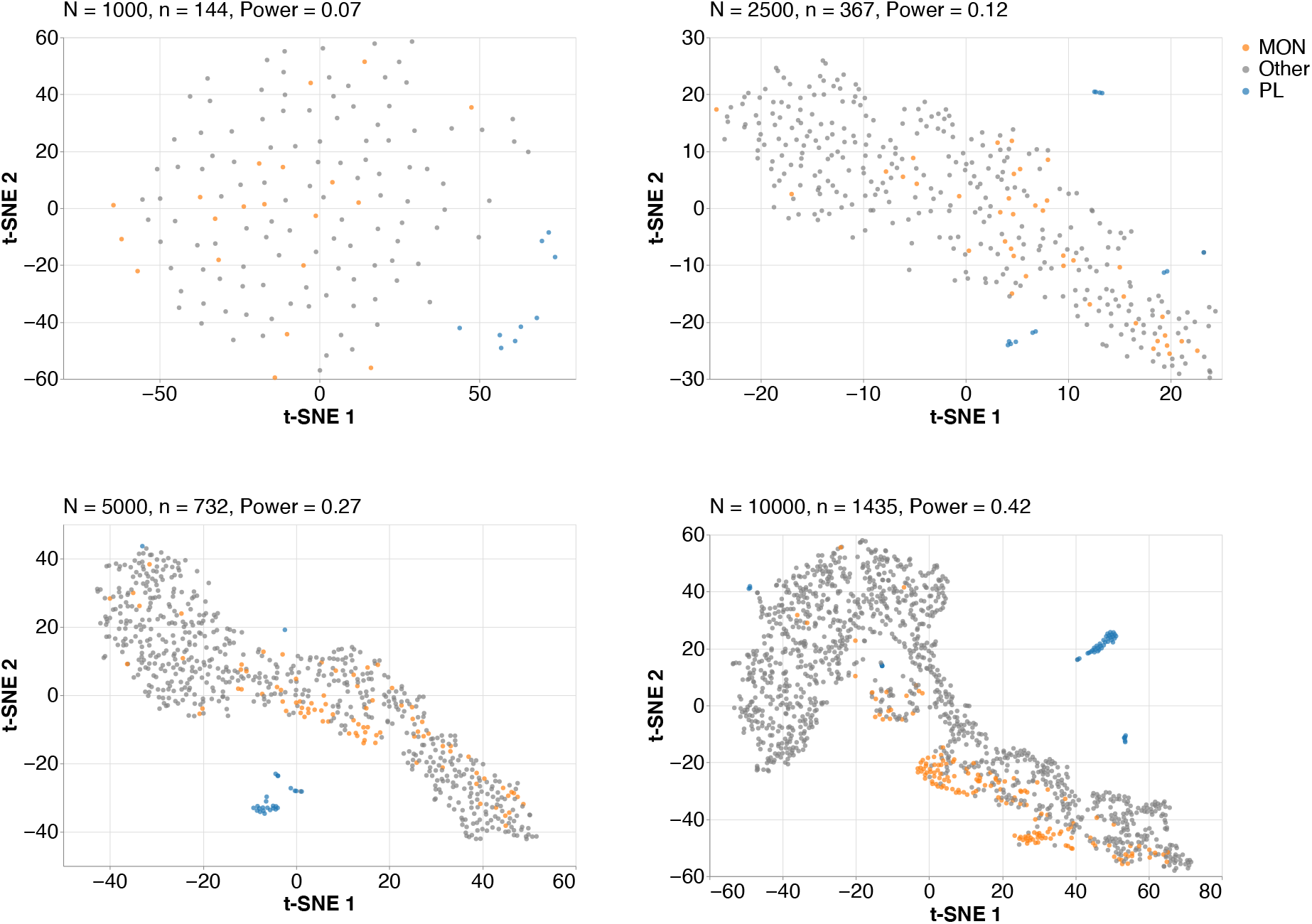
Latent representations of sub-clustering immune cells with varying samples sizes. Using samples of size N = (1000, 2500, 5000, 10000) from the KPMP data set [32], we extract immune cells and apply scVIDE to each of the resulting immune cell clusters to examine statistical power for sub-clustering. We show t-SNE representations of the lower-dimensional latent space of scVIDE and give the corresponding power estimates for each sample size n = (144, 367, 732, 1435) of the immune cells. Plasma (PL) cells are highlighted in blue and Monocytes (MON) are highlighted in orange.

## Notes

### Competing Interest Statement

The authors have declared no competing interest.

